# Ornithine Lipid is a Partial TLR4 Agonist and NLRP3 Activator

**DOI:** 10.1101/2022.01.28.477396

**Authors:** Malvina Pizzuto, Laura Hurtado-Navarro, Cristina Molina-Lopez, Jalal Soubhye, Michel Gelbcke, Silvia Rodriguez-Lopez, Jean-Marie Ruysschaert, Kate Schroder, Pablo Pelegrin

## Abstract

Myeloid cells recognise Gram-negative bacterial lipopolysaccharides (LPS). LPS recognition triggers inflammatory reactions through Toll-like receptor 4 (TLR4) and primes the cells for inflammasome activation. In phosphate-depleted environments, bacteria cannot produce LPS. Instead, they increase their synthesis of ornithine lipid (OL), which is constitutively present in some pathogenic Gram-negative and -positive bacteria but absent in commensals. OL is implicated in bacterial pathogenicity, but the mechanism is unclear. Using primary murine macrophages and human peripheral blood mononuclear cells, we identify OL as a partial TLR4 agonist and an NLRP3 inflammasome activator. For this, OL directly activates TLR4 and indirectly activates NLRP3 in a potassium-efflux-dependent manner. OL also upregulates the expression of NLRP3 and pro-IL-1β and induces cytokine secretion in primed and unprimed cells. By contrast, in the presence of LPS, OL functions as a partial TLR4 antagonist; LPS-induced TLR4 activation and inflammasome priming are inhibited by OL, leading to reduced TNF and IL-1β secretion. We thus suggest that in phosphate-depleted environments, OL replaces LPS bacterial immunogenicity while constitutively present OL may allow bacteria to escape immune surveillance.

## Introduction

In the realm of bacterial attacks on a host immune system, much is known about phospholipids like lipopolysaccharides (LPS) stimulating an inflammatory reaction in the host (Fitzgerald and Kagan, 2020; Violi *et al*., 2023). However, Gram-negative bacteria cannot produce such phospholipids if they encounter a low-phosphate environment, such as tropical forests and oceans(Bergkemper *et al*., 2016; Sosa *et al*., 2019); under these conditions Gram-negative bacteria produce amino-acid-containing lipids such as ornithine lipids (OLs) (Minnikin and Abdolrahimzadeh, 1974; Benning *et al*., 1993; López-Lara, Sohlenkamp and Geiger, 2003; VencesLJGuzmán *et al*., 2011). OLs are implicated in maintaining protein interactions, antibiotic resistance, membrane integrity and bacterial survival in those situations where phospholipids are not synthesised (Vences-Guzmán, Geiger and Sohlenkamp, 2012; Kim *et al*., 2018; Córdoba-Castro *et al*., 2021). However, OLs are not exclusively produced in phosphate-depleted medium but are also constitutively present at concentrations ranging from 2 to 26% in most Gram-negative bacteria (e.g., *Pseudomonas aeruginosa*, *Rhodobacter sphaeroides, Burkholderia sensu stricto* strains and *Brucella abortus*) (Rojas-Jiménez *et al*., 2005; Comerci *et al*., 2006; Sohlenkamp *et al*., 2007; Geiger *et al*., 2010; Barbosa *et al*., 2018) and some Gram-positive bacteria (e.g., *Mycobacterium tuberculosis* and *Streptomyces* (Laneelle *et al*., 1990) Interestingly, OLs are absent in avirulent strains of *Burkholderia* and *Mycobacterium,* and depleting OL production in pathogenic strains of *Burkholderia* reduced their ability to infect and cause mortality in eukaryotes (Córdoba-Castro *et al*., 2021), suggesting that OL is vital for bacteria pathogenicity. However, whether and how OL can interact with the eukaryote immune system is unclear (Córdoba-Castro *et al*., 2021). OLs purified from *B. pertussis*, *A. xylosoxidans* and *F. meningosepticum* induce tumor necrosis factor (TNF) and interleukin (IL)-6 and IL-1β pro-inflammatory cytokine production in macrophages (Kawai and Akagawa, 1989; Okemoto *et al*., 2008; Palacios-Chaves *et al*., 2011). Nevertheless, the function of OL in immunomodulation is not fully understood and is controversial since the ability of OL to induce cytokine secretion contrasts with other studies which show OLs as either inert or a suppressor of the bacterial LPS-induced immune response *in vivo* (Kawai *et al*., 1991; Palacios-Chaves *et al*., 2011). Such discrepancies may be due to the different bacterial origins and models used or LPS contamination in the bacterial-purified OLs, or alternatively, may reflect an unusual activity of OLs as a partial agonist of LPS receptors.

Given that OL’s ability to modulate host immune response may explain its role in bacterial pathogenicity in either the presence or absence of LPS, in this study, we investigated the capacity of synthetic OL to activate or inhibit the immune response in primary murine and human myeloid cells, and the receptors involved.

Synthetic ionisable lipids with structural similarity to OL activate innate immune receptors in myeloid cells, such as Toll-like receptors (TLR) 2 and 4 and the nucleotide-binding domain and leucine-rich repeat-containing pyrin protein 3 (NLRP3) inflammasome, inducing pro-inflammatory cytokines such as those reported to be induced by OLs (e.g., IL-6, TNF and IL-1β) (Lonez *et al*., 2015; Pizzuto *et al*., 2018). We thus hypothesised that activation/inhibition of these receptors by OLs could be responsible for the observed OL-induced/inhibited cytokine secretion. TLRs are transmembrane proteins that detect specific microbial signatures (Barton and Medzhitov, 2003; Paramo *et al*., 2013; Fitzgerald and Kagan, 2020). For example, TLR4 recognises Gram-negative bacterial lipids such as LPS, while TLR2 recognises lipopeptides from Gram-positive bacteria and their synthetic derivatives such as Pam_3_CSK_4_ (Kang *et al*., 2009; Park *et al*., 2009). When TLRs bind to their ligand, they activate several transcription factors, including the nuclear transcription factor (NF)-κB that induces the production and secretion of inflammatory cytokines such as TNF and IL-6. NF-κB activation also induces the expression of inflammasome pathway proteins, including NLRP3 and pro-IL-1β. Thus, treating cells with a TLR agonist followed by an NLRP3 stimulus activates the NLRP3 inflammasome, a large signalling complex formed by NLRP3, the apoptosis-associated speck-like protein containing a caspase recruitment domain (ASC) protein and active caspase-1. Active caspase-1 cleaves pro-IL-1β and GSDMD to their active pro-inflammatory forms of IL-1β and Nt-GSDMD. Nt-GSDMD then forms pores in the plasma membrane, inducing IL-1β secretion and a lytic form of cell death called pyroptosis, which results in the release of intracellular content such as lactate dehydrogenase (LDH) and pro-inflammatory damage-associated molecular patterns (DAMPs). NLRP3 can be activated by a decrease in K^+^ intracellular concentrations induced either by ATP or toxins such as nigericin, among others, and by GSDMD-induced plasma membrane permeabilisation via caspase-11 activation by cytoplasmic LPS (non-canonical inflammasome) (Schroder and Tschopp, 2010; Kayagaki *et al*., 2011; Hagar *et al*., 2013; Broz, Pelegrín and Shao, 2019). In the present work, we sought to determine whether synthetic OL could activate these innate immune pathways in human and murine myeloid cells. We identified OL as an activator of the canonical NLRP3 inflammasome and a partial agonist of TLR4. Thus, when administered alone, OL could induce both priming and activation of NLRP3 to induce the secretion of TNF, IL-1β and other cytokines. When administered in combination with LPS, OL inhibited LPS-induced TLR4 activation. This study gives new insight into the immunomodulatory properties of OL, and provides a molecular explanation for the apparent discrepancies in the current literature.

## Materials and Methods

### Reagents

Ultrapure lipopolysaccharide (LPS) from *E. coli* 0111:B4, Pam_3_CSK_4_, mouse TLR2 neutralising antibody monoclonal mouse IgG2a (C9A12) and the control isotype mouse IgG2a were from InvivoGen. Mouse TLR4/MD2 complex neutralising antibody (monoclonal rat IgGk clone MTS510) and control isotype rat IgGk were from eBioscience™ Invitrogen. FuGENE® was from Promega, while cholera toxin B (CTB) was from Sigma. C14:0 ornithine lipid (OL) was synthesised as described in the Extended View file.

### OL liposome preparation

CHCl_3_ solutions of 1 mg/mL OL were prepared, and lipid films were prepared by solvent evaporation under a nitrogen stream before being dried overnight and kept at −20°C. Before each experiment, liposomes were freshly formed by resuspending lipid films into filtered HEPES 10 mM (BioWhittaker), heating for 20 minutes at 70°C, and then sonicating for 5 min in a sonicator bath (Sonitech).

### LPS/Fugene**®** and CTB/LPS complex preparation for intracellular delivery of LPS

LPS (InvivoGen) was first mixed with Fugene® (Promega) in a small volume of Opti-modified Eagle’s medium (MEM)™ reduced-serum medium (Invitrogen) that was previously heated to 37°C. The mix was vortexed and then incubated for 15 minutes at room temperature to allow the formation of the complex, and finally, more Opti-MEM™ was added to reach a final concentration of 0.5% Fugene and 1 μg/mL LPS (LPS/Fu). Alternatively, a mix of LPS and 20 ug/mL of cholera toxin B (CTB) from Sigma was made in Opti-MEM™ before vortexing and then incubating for 5 min at room temperature to allow the formation of the complex (CTB/LPS).

### HEK-Blue™ cell line culture and treatments

HEK-Blue™ mTLR4 cells (InvivoGen) were cultured in Dulbecco’s modified Eagle’s medium (DMEM) F12 (Biowest) supplemented with 10% heat-inactivated Fetal Bovine Serum Premium (Biowest), 1% L-glutamine (Lonza), and from the second passage, 0.1 mg/mL of Normocin™ (InvivoGen) and 1X HEK-Blue™ selection antibiotic (InvivoGen). Cells were maintained at 37°C in a 5% CO_2_ atmosphere and tested regularly for mycoplasma contamination with a MycoProbe Mycoplasma Detection Kit following the manufacturer’s instructions (R&D Systems). Cell cultures were passaged no more than 15 times, to avoid divergence from the parent line. For experiments, HEK-Blue™ mTLR4 cells were seeded at the rate of 70,000 cells and 100 μL of culture medium per well in 96-well plates (Greiner Bio-One). After 24 hours, and with cells at 80% confluence, the supernatant was discarded and replaced with either 23 μL of control solution (HEPES and water) alone or OL or LPS diluted in the control solution and 77 μL of HEK-Blue™ Detection medium (InvivoGen) (cell culture medium for real-time detection of secreted alkaline phosphatase, SEAP). Following the manufacturer’s instructions, TLR4-dependent NF-κB activation was measured by the real-time detection of SEAP, based on the absorbance of the HEK-Blue™ detection medium at 620 nm, measured using a BioTek Synergy HT Microplate Readers at 37°C after 4, 22, and 29 hours of incubation in a 37°C/5% CO_2_ atmosphere and reported as fold increase with respect to the control condition. Since the highest fold increase induced by both LPS and OL was at 22 hours, we chose to focus experimental end points on this time point.

### Differentiation of bone marrow-derived macrophages (BMDMs)

Experiments conducted at the Biomedical Research Institute of Murcia used C57 BL/6J, *Nlrp3*^−/−^, *Casp-1/11*^−/−^ or Pycard(*Asc)*^−/−^male and female mice between 8 and 13 weeks of age and bred under specific pathogen-free (SPF) conditions, in accordance with the *Hospital Clínico Universitario Virgen Arrixaca* animal experimentation guidelines, and the Spanish national (RD 1201/2005 and Law 32/2007) and EU (86/609/EEC and 2010/63/EU) legislation. Accordingly, no specific procedure approval is needed when animals are sacrificed to obtain biological material. Bone marrow obtained from the leg bones of mice euthanised by CO_2_ inhalation was used to differentiate bone marrow-derived macrophages (BMDMs) that were used at day 7 of differentiation. Briefly, the bone marrow was flushed from the bone cavity and resuspended in differentiation medium (DMEM media, with L-glutamine, without sodium pyruvate (Biowest) supplemented with 10% heat-inactivated fetal bovine serum premium (Biowest), 2 mM glutamine (Lonza), 50 U/mL penicillin, 50 μg/mL streptomycin (PEN-STREP, Lonza), and 20% of supernatant from the L929 cultures). The bone marrow cell suspension was maintained in Petri dishes in a 37°C/5% CO_2_ atmosphere. After two days, the differentiation medium was supplemented, and cells were maintained for four extra days.

Alternatively, experiments at the Institute for Molecular Bioscience (University of Queensland) used C57BL/6, *Tlr4*^−/−^, *Casp-11*^−/−^ and *Nlrp3*^−/−^ male and female mice between 6 and 14 weeks of age and bred under specific pathogen-free (SPF) facilities at the University of Queensland. All protocols involving mice were approved by the University of Queensland Animal Ethics Committee and we have complied with all relevant ethical regulations. Murine macrophages were differentiated from bone marrow and were used for experiments on day 7 of differentiation. Briefly, the bone marrow was flushed from the bone cavity filtered, spun down at 400 g and resuspended in differentiation medium consisting of RPMI 1640 medium (Life Technologies) supplemented with 10% heat-inactivated fetal calf serum (FCS), 2□mM GlutaMAX (Life Technologies), 50□U per ml penicillin–streptomycin (Life Technologies) and 150□ng□ml^−1^ recombinant human macrophage colony-stimulating factor (CSF-1 (endotoxin-free, expressed and purified by the University of Queensland Protein Expression Facility)). The bone marrow cell suspension was maintained in Petri dishes in a 37°C/5% CO_2_ atmosphere. After five days, the differentiation medium was supplemented, and cells were maintained for one extra day.

### BMDM stimulation

BMDMs differentiated for six days were washed and detached from their Petri dishes using Dulbecco’s phosphate-buffered saline (PBS) (Thermo Fisher Scientific). The cells were then counted, centrifuged for 5 min at 500 g (Centrifuge 3-18KS, Sigma), then resuspended in full media to a concentration of 1 × 10^6^ cells/mL and distributed in 96-well plates (100 μL/well), 24-well plates (500 μL/well), or 6-well plates (2 mL/well) (Greiner Bio-One).

For experiments involving overnight incubations, after 3 hours, 10, 50, or 200 μL of the medium was added, either alone or supplemented with Pam_3_CSK_4_ 1 μg/mL. After a further 4 hours, supernatants were collected, and the cells were washed with PBS. Control solution (HEPES and Opti-MEM™), OL, LPS/CTB, or LPS/Fu diluted in control solution were added and cells incubated for 18 hours. Alternatively, control solution (HEPES and Opti-MEM™) or OL was added, and cells were incubated for 17 hours, followed by 1 hour incubation with nigericin. For experiment involving BMDMs exposed to stimulants for 1 or 4 hours, BMDMs were differentiated for six days and replated as above then left for 24 hours (rather than 3 hours) in differentiation medium, which was then discarded and control solution (HEPES and Opti-MEM™), OL, Pam_3_CSK_4_ or LPS diluted in Opti-MEM™ in the absence or presence of neutralising antibodies or antibodies controls, were added and cells incubated for 4 h. Alternatively, Opti-MEM™ alone or with nigericin were added and cells incubated for 1 hour. The figure legends indicate doses of stimuli, BMDM phenotypes, and combinations of OL with other stimuli. After incubation, the supernatants were collected, centrifuged at 600 g, and assayed for cytokine and LDH release, while cell lysates were assayed for protein or mRNA expression.

### Isolation of peripheral blood mononuclear cells (PBMC)

Whole peripheral blood samples were collected in EDTA anticoagulation tubes from healthy donors, as approved by the Institutional Review Board of the *Hospital Clínico Universitario Virgen de la Arrixaca*. Informed consent was obtained from all individuals enrolled in the study following the principles set out in the WMA Declaration of Helsinki, and samples were stored in the Biobanco en Red de la Región de Murcia (PT13/0010/0018) integrated in the Spanish National Biobanks Network (B.000859). PBMC were obtained by Ficoll gradient centrifugation in Histopaque-1077 (Sigma-Aldrich) using SepMate™ isolation tubes from STEMCELL™ and following the manufacturer’s instructions. Isolated PBMC were resuspended in RPMI 1640 (Sigma-Aldrich) supplemented with 10% heat-inactivated FBS premium (Biowest) and maintained in a 37°C/5% CO_2_ atmosphere.

### PBMC treatments

Suspensions of freshly-isolated PBMC were distributed in round-bottom 96-well plates (50 μL, 100,000 cells per well) with 50 μL of RPMI 1640 and control solutions (HEPES and water) with or without OL in the absence or presence of MCC950 10 μM (MedChemExpress). All cells were maintained in a 37°C/5% CO_2_ atmosphere for 4 or 18 hours before the supernatants were collected, centrifuged at 600 g, and then assayed for cytokine secretion.

### Cytokine assays

Murine TNF and IL-1β were quantified in cell supernatants using the ELISA™ Kit from Invitrogen and human IL-1β was similarly quantified with the Instant ELISA™ Kit from Invitrogen, following the manufacturer’s instructions. Absorbance was read with a BioTek Synergy HT Microplate Reader. Alternatively, murine TNF and IL-1β were quantified in cell supernatants using the DuoSet® ELISA Kit from RnD system following the manufacturer’s instructions. Absorbance was read with a Tecan Microplate Reader. All other cytokines were quantified in cell supernatants using a custom Multiplex Kit procartaplex from Thermo Fisher Scientific and a Luminex MAGPIX System (Millipore), as per the manufacturer’s instructions. Cytokine amounts were reported as ng or pg per million cells to standardise the difference in cell amount/volume of media ratio between the different plate layouts.

### LDH assay

For the experiments conducted at the Biomedical Research Institute of Murcia, LDH activity was quantified in the cell lysate of untreated cells and in all cell supernatants using the Cytotoxicity Detection Kit (LDH) from Merck, following the manufacturer’s instructions. Absorbances at 492 and 620 nm were measured with a BioTek Synergy HT Microplate Reader every minute for 20 min, with the slopes of increase in absorbance calculated with respect to time, and background values subtracted from the value of each supernatant reported as percentages of the sum of the value measured in the supernatant and the lysate of the untreated condition (total LDH). The lysate was obtained as follows: cells were lysed with 2% Triton lysis buffer comprising 150 mM NaCl (Immobilon), 2% Triton X-100 (Sigma-Aldrich), and 50 mM Tris-HCl pH8 (Sigma-Aldrich) supplemented with 100 µL/mL of protease inhibitor (Sigma-Aldrich). Cells were scraped in cold lysis buffer on ice. Lysates were then incubated for 30 minutes on ice with a vortex every 10 minutes, before centrifugation for 10 minutes at 13,000 g in a microcentrifuge (1-14K, Sigma) to remove the pellets containing cell debris.

Alternatively, LDH activity was quantified in all cell supernatants using the Cytox96 non-radioactive cytotoxicity assay (Promega). Absorbances at 492 and 620 nm were measured with a Tecan Microplate Reader and reported as percentages of the value measured in the supernatant of cells treated with 1% triton for 10 minutes (total LDH).

### Quantitative PCR

BMDM treated in 6-well plates were washed twice with PBS before total RNA purification using the RNeasy kit (Qiagen) following the manufacturer’s recommendations, with RNA quantified using a nanodrop 2000 (ThermoFisher). Reverse transcription was performed using iScriptTM cDNA Synthesis kits (BioRad) according to the manufacturer’s instructions, and quantitative PCR (qPCR) was performed in an iQTM 5 Real Time-PCR Detection System (BioRad) with a SYBR Green mix (Takara and predesigned primers obtained from Sigma-Aldrich (KiCqStart® Primers). The presented relative gene expression levels were calculated using the 2^−ΔΔCT^ method (Livak and Schmittgen, 2001) with Glyceraldehyde 3-phosphate dehydrogenase (GAPDH) as endogenous controls.

### Endotoxin test

LPS was quantified in OL liposomes using the Pierce™ Chromogenic Endotoxin Quant Kit (Thermo Fisher), following the manufacturer’s instructions.

### Statistical analysis

Each graphics symbol presented represents the mean value of biological triplicates from an independent experiment, while each bar represents the mean value, and the error bars represent the standard error of the mean (SEM) of three or more independent experiments, as indicated in the figure. A two-tailed unpaired, paired, or ratio *t*-test was used for two-group comparisons, as indicated in the figure. Comparisons of multiple groups were analysed by one-way analysis of variance (ANOVA) with Dunnet’s or Tukey’s multiple-comparison test, as indicated in the figure. A significant *p*-value is indicated as \**p*<0.05; and *p*>0.05 as not significant (ns or not indicated). Nonlinear regression analysis was carried out using GraphPad Prism 9 software software (Graph-Pad Software, Inc.). Prism 9 was also used to generate graphs, calculate SEM and perform statistical analysis.

## Results

### OL is a partial agonist of TLR4

Previous studies show that OL induce pro-inflammatory cytokine secretion, while others suggest OL inhibit LPS-dependent pro-inflammatory reactions. We hypothesise that such a discrepancy occurs because OL may be a partial agonist of the LPS receptor, Toll-like receptor 4 (TLR4). Indeed, partial agonists weakly activate their receptors when administered alone but inhibit the activation of those same receptors by strong agonists (Kenakin, 2017). To investigate this hypothesis, we synthesised an Ornithine Lipid with 14 carbon atom chain length (Fig 1 A and EV 1) and compared its immunogenicity to the immunogenicity of LPS. At first, we monitored TLR4 responses using HEK cells stably transfected with TLR4 plus a secreted phosphatase (SEAP) as an NF-KB reporter (HEK-TLR4). We then measured the NF-κB activation induced by increasing concentrations of OL alone (Fig 1 B) or by increasing concentrations of LPS in the absence or presence of a fixed OL concentration (Fig 1 C). The resulting dose-response curves showed that when administered alone, OL induces dose-dependent NF-κB activity in HEK-TLR4 cells that is approximately two times lower than that induced by LPS alone (Fig 1 B and C). Intriguingly, when administered alongside LPS, OL inhibited LPS-induced TLR4 signalling for LPS concentrations lower than 50 ng/mL (Fig 1 C). Increasing LPS concentrations in the presence of OL (dashed line Fig 1 C) could eventually reach the same NF-κB activation of LPS alone (solid line, Fig 1 C), which strongly suggests that OL competes with LPS for the same binding site on TLR4. Together, these data obtained in the reductionistic system of HEK-TLR4 cells indicate that OL is a partial TLR4 agonist.

**Figure 1:**
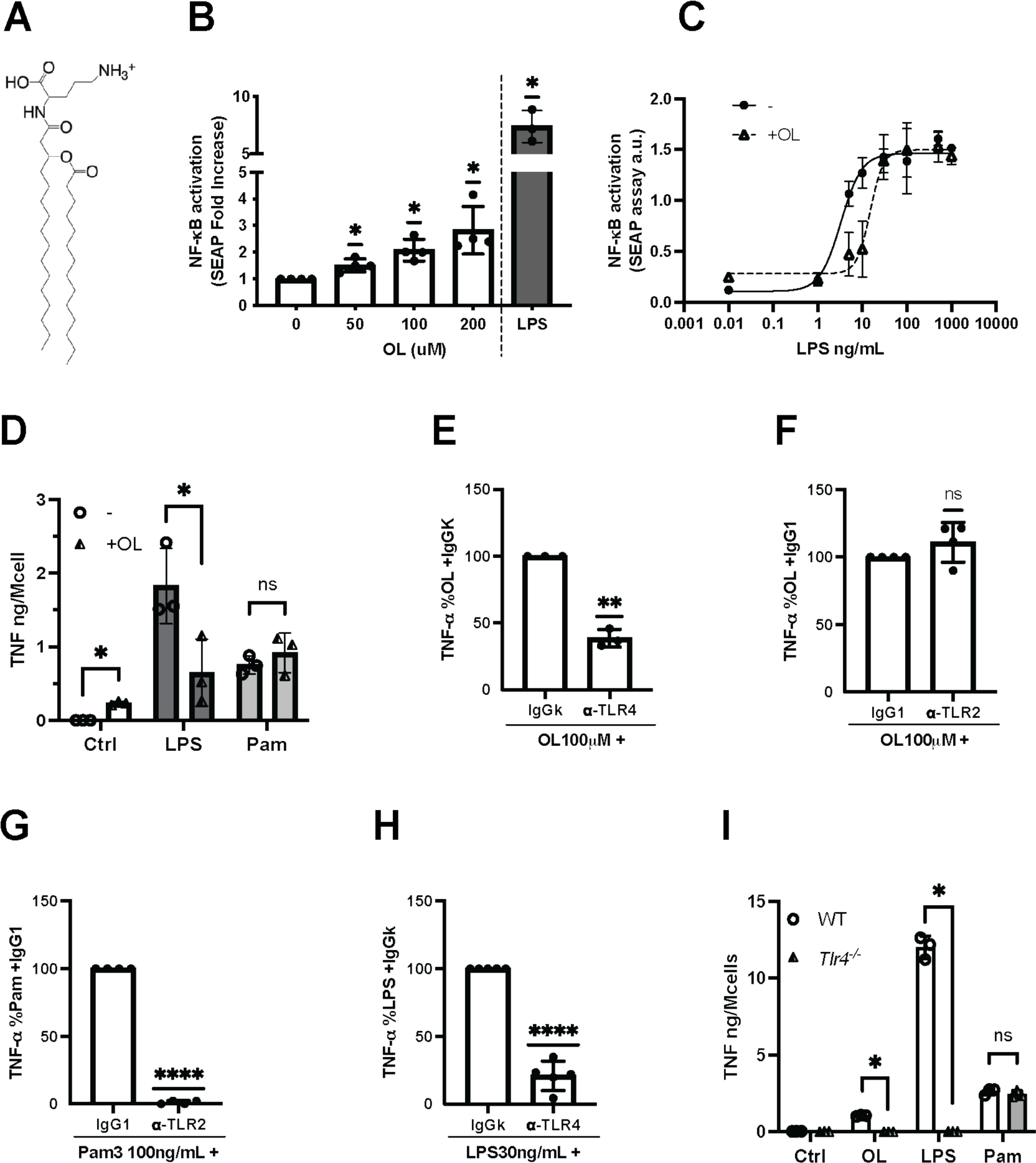
OL is a par/al TLR4 agonist. **A** HEKBlue-mTLR4 cells were incubated for 22 hours with medium and control solution (0), increasing concentrations of OL (50 to 200 μM), or LPS 100 ng/mL (grey bar). TLR4-dependent NF-κB activation was quantified by secreted alkaline phosphatase (SEAP) assay and reported here as a fold increase with respect to the control condition. **B** HEKBlue-mTLR4 cells were incubated for 4 hours with increasing concentrations of LPS (0.01 to 1000 ng/mL) in the absence (circles, solid line) or presence of OL 100 μM (triangles, dashed line). TLR4-dependent NF-κB activation was quantified by secreted alkaline phosphatase (SEAP) assay and reported here as an arbitrary unit. Nonlinear regression analysis was carried out using GraphPad Prism 9 soXware. **C** BMDM from wild-type mice were incubated for 4 hours with medium (white bars), LPS 100 ng/ml (dark grey bars), or Pam_3_CSK_4_ 1 μg/mL (light grey bars) in the absence (circles) or presence of OL 100 μM (triangles). TNF was quantified in collected supernatants by ELISA. **D-G** BMDM from wild-type mice were incubated for 4 hours with OL 100 μM (D and E), LPS 30 ng/ml (F), Pam_3_CSK_4_ 100 ng/mL (G), in the presence of 5 μg/mL of antibodies blocking TLR4 (D and F) or TLR2 (E and G), or their respective isotype control antibodies IgGk (D and F) or IgG1 (E and G). TNF was quantified in collected supernatants by ELISA and reported here as the percentage of the secretion measured in the presence of both the activator and isotype control antibody. **H** BMDM from wild-type mice (circles, white bars) or from *Tlr4_-_*_/-_ mice (triangles, grey bars) were incubated for 4 hours with LPS 100 ng/ml, Pam_3_CSK_4_ 500 ng/mL, or OL 200 μM. TNF was quantified in collected supernatants by ELISA. Data information: Bars are the mean of three or more independent experiments represented by symbols (n ti 3 to 5) ± SEM. Statistical analysis: A, D, E, F, and G one-sample t-test. C and H Ordinary one-way ANOVA Tukey’s multiple comparisons test (paired). Significant difference for p<0.05 (*). Only comparisons of interest are shown.

Next, we investigated the ability of OL to induce and inhibit TLR4-dependent inflammatory cytokine secretions in myeloid cells. First, we incubated BMDMs with OL in the presence or absence of LPS or the TLR2 agonist Pam_3_CSK_4_, with the latter included to investigate OL specificity towards TLR4. As shown in Figure 1 D, OL induced TNF secretion in BMDMs (white bar) but also inhibited TNF release induced by LPS (dark grey bars, circles vs triangles). In addition, OL did not inhibit TNF induced by Pam_3_CSK_4_ (light grey bars, circles vs triangles). Thus, when administered alone, OL induces pro-inflammatory pathways in murine macrophages. When co-administered with LPS, OL specifically inhibits LPS-induced TLR4 signalling.

To next test whether OL-induced TNF secretion could reflect OL-triggered TLR4 activation in macrophages, we measured OL-induced TNF secretion in BMDMs pre-incubated with either antibodies blocking TLR4 or TLR2 or with their respective antibody controls (IgG1 or IgGk). We observed that OL-induced TNF secretion was prevented by antibodies blocking TLR4 (Fig. 1 E) but not TLR2 (Fig. 1 F), demonstrating that the OL-induced TNF secretion in BMDMs requires TLR4 but not TLR2. BMDMs stimulated with either the TLR4 agonist LPS or the TLR2 activator Pam_3_CSK_4_ served as positive controls, and Pam_3_CSK_4_- and LPS-induced TNF secretion were prevented by TLR2 and TLR4 neutralisation, respectively (Fig. 1 G and H). To further confirm the ability of OL to activate TLR4, we compared OL-induced TNF secretion in BMDMs derived from WT (white bars, circles) versus *Tlr4*^-/-^ mice (grey bars, triangles). BMDMs stimulated with either the TLR4 agonist LPS or the TLR2 activator Pam_3_CSK_4_ served as positive controls. Pam_3_CSK_4_ induced TNF secretion in *Tlr4*^-/-^ BMDM as anticipated, while OL- and LPS-induced TNF secretion was ablated in *Tlr4*^-/-^ BMDM (Fig. 1 I), confirming that OL induces TNF secretion in primary macrophages through TLR4 activation.

Macrophages are exquisitely sensitive to LPS, including the trace levels of contaminating LPS found in some cell culture reagents purified from bacteria. Therefore, we used synthetic OL in all experiments, and verified that this reagent was not contaminated by LPS (Fig. EV2). Collectively, these observations demonstrate that OL alone can weakly activate TLR4.

### OL induces a broad spectrum of pro-inflammatory cytokines in human and murine myeloid cells

To further characterise the inflammatory responses caused by OL, we incubated BMDMs with OL concentrations ranging from 25 to 200 µM and measured TNF secretion by ELISA. As shown in Figure 2 A, the TNF secretion induced by OL was dose-dependent. Substantial TNF secretion was observed with as little as 50 µM OL and this was further elevated at 100 and 200 µM OL. Therefore, we chose to carry out multiplex assays with supernatants of BMDMs treated with 100 and 200 µM OL to measure other TLR4-dependent cytokines such as IL-6, CXCL10, IL-2 and IL-10. These analyses revealed that OL induced a significant and dose-dependent increase in the secretion of all tested cytokines (Figure 2 B to E), indicating that OL induces a broad cytokine secretion program. We next sought to understand whether OL shows similar immunogenicity in human cells. We incubated human peripheral blood mononuclear cells (PBMC) with 100 or 200 µM OL and measured TNF secretion by ELISA after 4 and 18 hours. After 4 hours, OL could induce only a weak but dose-dependent TNF secretion in PBMC (Fig 2 F), which was stronger after 18 hours of incubation with OL (Fig 2 G). Thus, we conclude that OL also induces pro-inflammatory cytokines in human cells, although with a slower kinetic.

**Figure 2:**
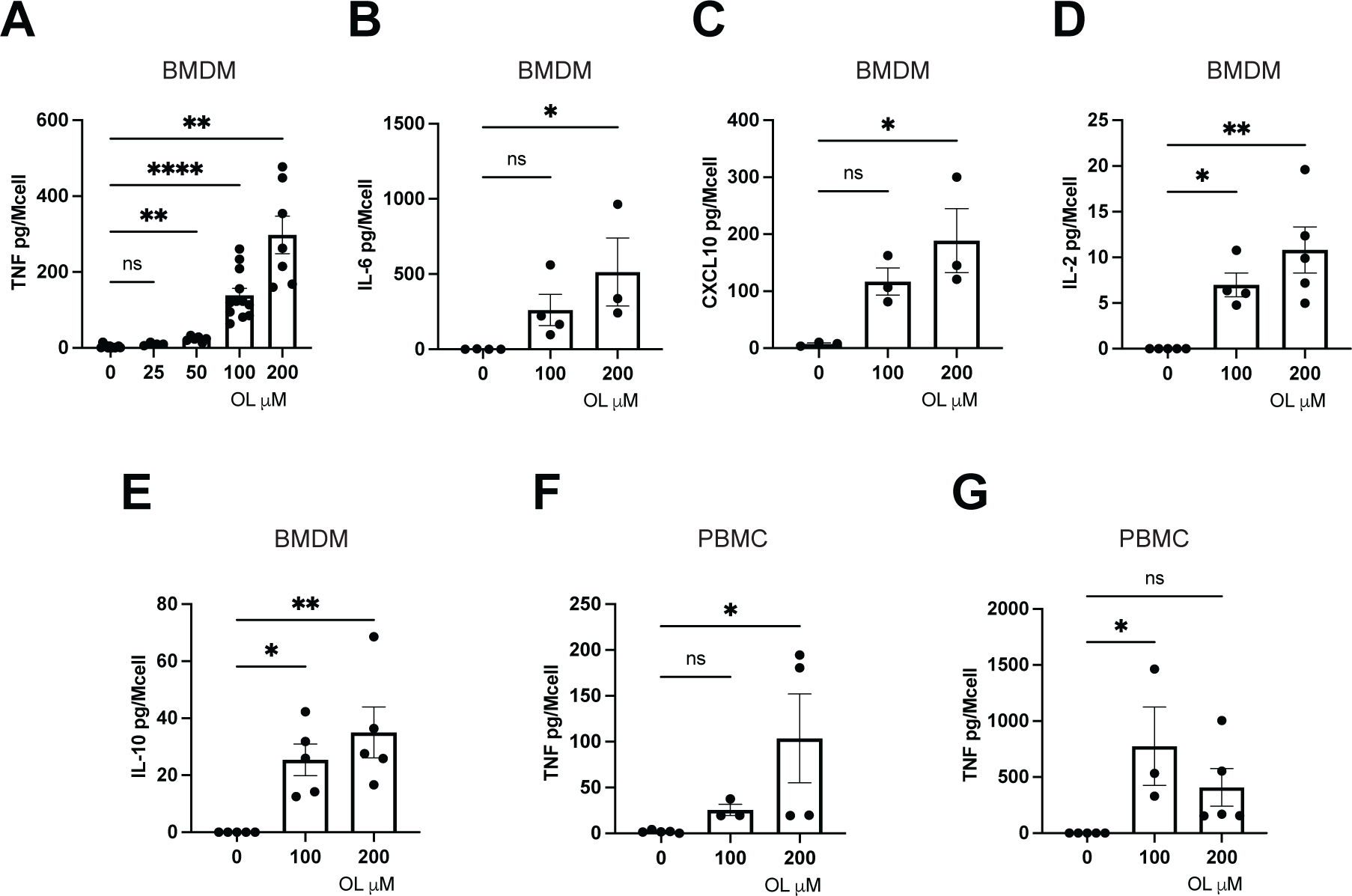
Characteriza/on of OL-induced pro-inflammatory response in human and murine myeloid cells. **A to E** BMDM from wild-type mice were incubated for 4 hours without (0) or with increasing concentrations of OL (from 25 to 200 µM). TNF (**A**) was quantified in supernatants by ELISA. IL-6 (**B**), CXCL10 (**C**), IL-2 (**D**), and IL-10 (**E**) were quantified by Luminex assay. **F to G** Human PBMCs from healthy donors were incubated for 4 (**F**) or 18 (**G**) hours without (0) or with increasing concentrations of OL (from 100 to 200 µM). TNF was quantified in supernatants by ELISA. Data information: Bars are the mean of three or more independent experiments represented by symbols (n ti 3 to 5) ± SEM. Statistical analysis: Ordinary one-way ANOVA Dunnej’s multiple comparisons test (paired), significant difference for p<0.05 (*). Only comparisons of interest are shown.

### OL downregulates LPS-induced non-canonical inflammasome signalling by specifically inhibiting TLR4

Based on our finding that OL regulates the extracellular LPS receptor, TLR4, we wondered whether it could also modulate the intracellular LPS receptor, caspase-11 (casp-11). When LPS is delivered intracellularly with transfection agents such as Fugene® (LPS/Fu) or Cholera Toxin B (LPS/CTB), it activates murine casp-11 and induces the non-canonical inflammasome pathway. This cascade results in NLRP3-independent pyroptosis and the resultant release of intracellular contents, including lactate dehydrogenase (LDH), and also induces a second wave of inflammasome signalling (NLRP3-ASC-casp-1) to trigger IL-1β maturation and release. Therefore, to test whether OL could specifically activate or inhibit casp-11, we primed BMDMs with the TLR2 agonist Pam_3_CSK_4_to upregulate expression of NLRP3 and pro-IL-1β and co-incubated primed BMDMs with OL in the absence or presence of activators of the noncanonical inflammasome (LPS/CTB) or the canonical NLRP3 inflammasome (nigericin). OL did not decrease LPS/CTB-induced LDH release (Fig. 3 A), suggesting that OL does not inhibit LPS-induced caspase-11 activity and resultant pyroptosis. OL did, however, suppress IL-1β secretion stimulated by intracellular LPS but not by nigericin (Fig. 3 B), suggesting that OL may suppress the abundance of pro-IL-1β available to inflammasomes, perhaps by suppressing LPS-TLR4 signalling. To confirm this hypothesis, we next tested whether the observed inhibition of intracellular LPS-induced IL-1β release could be reproduced in BMDMs from *Tlr4^-/-^* mice. Indeed, OL failed to inhibit LPS/CTB-induced IL-1β release in *Tlr4^-/-^* macrophages (Fig. 3 C), confirming that the suppressive effect of OL on LPS-induced IL-1β production requires TLR4. Thus, OL limits the abundance of pro-IL-1β available to inflammasomes by suppressing LPS/TLR4-induced pro-IL1β expression.

**Figure 3:**
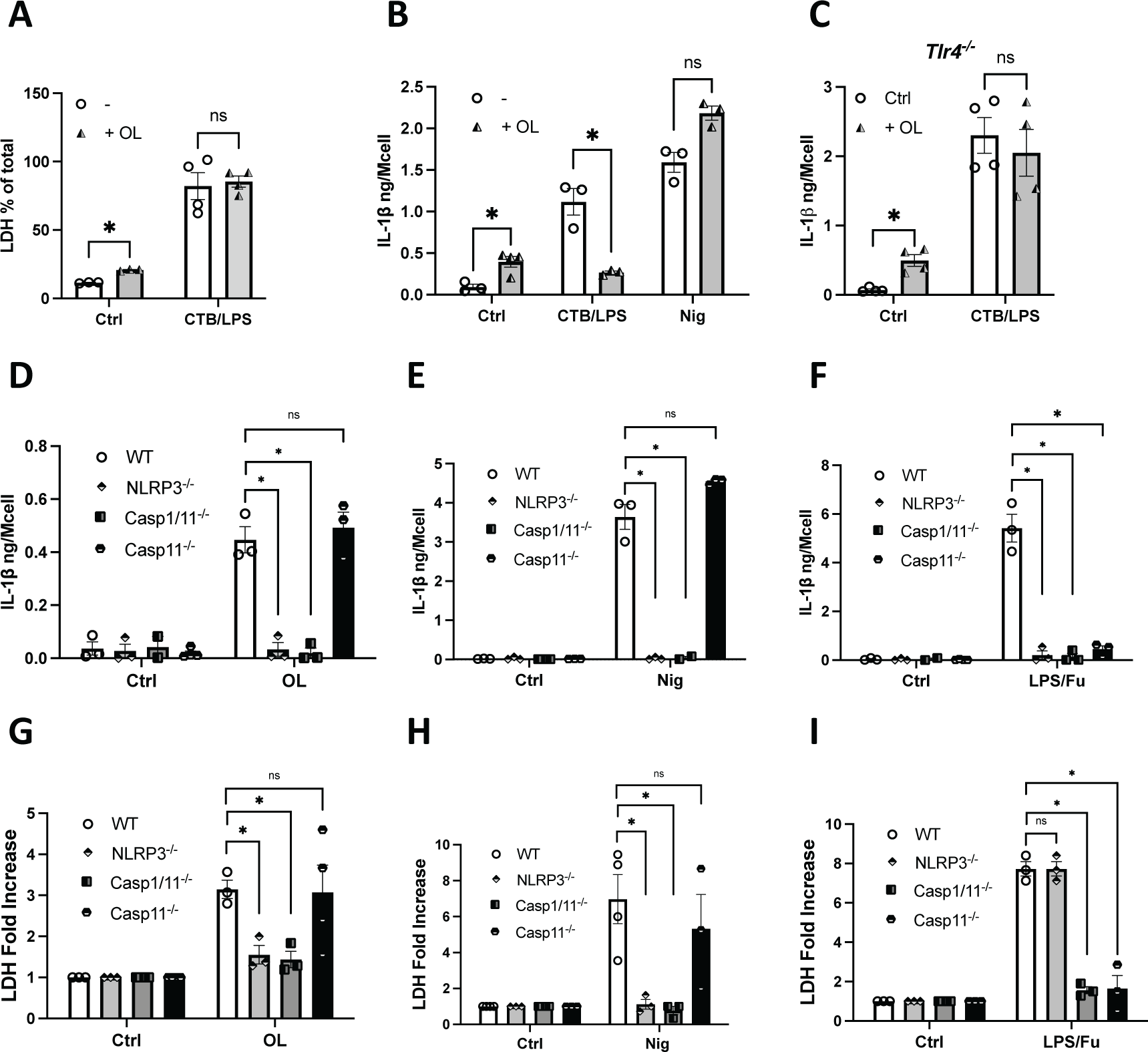
OL downregulates specifically LPS-induced non-canonical inflammasome signaling through TLR4 inhibi/on, while ac/va/ng the canonical NLRP3 inflammasome. **A to C** BMDM from wild type (A and B) or *Tlr4*^-/-^ (C) mice were incubated for 4 hours with Pam_3_CSK_4_ (1 µg/mL), then washed and incubated for 18 hours with medium (Ctrl) or CTB/LPS 20/2 µg/mL in the absence (-) or presence of OL 100 µM, or for 17 hours with medium (Ctrl) or OL 100 µM followed by 1 hour with medium (ctrl) or nigericin 10 µM. LDH activity was measured in fresh supernatant by cytotoxicity LDH assay and reported here as the percentage of LDH activity measured in total cell lysate (A). IL-1β was quantified in cell supernatants by ELISA (B and C). **D to I** BMDM from wild type, *Nlrp3*^-/-^, *Casp1/11*^-/-^ or *Casp11*^-/-^ mice were incubated for 4 hours with Pam_3_CSK_4_ (1 µg/mL), then washed and incubated for 18 hours with medium or OL 100 µM (D and G), with medium or LPS/Fu 1 µgmL^-1^/0,5% (F and I), or incubated for 1 hour with medium or nigericin 10 µM (H and I). IL-1β was quantified in cell supernatants by ELISA (D, F, and H). LDH activity was measured in fresh supernatant by cytotoxicity LDH assay and reported here as fold increase with respect to the phenotype ctrl (E, G, and I). **Data information:** Bars are the mean of three or more independent experiments represented by symbols (n ti 3 to 4) ± SEM. **Statistical analysis:** A to C and G to I unpaired t-test. D to F one-way ANOVA Tukey’s multiple comparisons test (paired), significant difference for p<0.05 (*). Only comparisons of interest are shown.

### OL activates the canonical NLRP3 inflammasome

Intriguingly, when OL was administered to cells without LPS or nigericin, OL induced the release of both IL-1β (Fig. 3 A) and LDH (Fig. 3 B). This suggested an additional role for OL as an inflammasome activator. To test this hypothesis, we primed BMDMs derived from wild-type, NLRP3 (*Nlrp3*^-/-^), caspase-1/11 (*Casp1/11*^-/-^) or caspase-11 (*Casp11*^-/-^)-deficient mice with the TLR2 activator Pam_3_CSK_4_ for 4 hours prior to incubation with OL, and monitored OL-induced IL-1β and LDH release, using nigericin and intracellular LPS (LPS/Fu) as positive controls for the canonical and non-canonical pathway, respectively. As expected for a non-canonical inflammasome activator, LPS/Fu induced IL-1β secretion in wild-type BMDM but not *Nlrp3^-/-^, Casp-1/11*^-/-^, and *Casp-11*^-/-^ BMDM (Fig. 3 F), whereas LPS/Fu induced LDH release wild-type and *Nlrp3*^-/-^ BMDM but not *Casp-1/11*^-/-^ and *Casp-11*^-/-^ BMDM (Fig. 3 I). OL induced IL-1β and LDH release in primed wild-type and *casp-11^-/-^* BMDM but not *Nlrp3*^-/-^ or *Casp-1/11*^-/-^ BMDM, mirroring the profile of the canonical NLRP3 activator, nigericin (Fig. 3 D, E, G and H). These data demonstrate that OL is a canonical NLRP3/casp-1 activator that does not activate casp-11.

### OL activates TLR4 and the NLRP3 inflammasome independently of one another

TLR4 and NLRP3 pathways in myeloid cells are interconnected, with TLR4 activation enhancing NLRP3 signalling by increasing the expression of inflammasome components. Moreover, in certain conditions, NLRP3 can be activated downstream of TLR4, such as in LPS-treated monocytes (Gaidt *et al*., 2016). We, therefore, investigated whether OL can independently induce NLRP3 and TLR4 signalling pathways. As shown in Figure 4, OL can induce TNF secretion in BMDMs from mice unable to signal via the NLRP3 inflammasome (*Asc*^-/-^) (Fig. 4 A), and OL-induced IL-1β and LDH release were maintained in BMDMs from *Tlr4*^-/-^ mice primed with Pam_3_CSK_4_ (Fig. 4 B and C). These findings demonstrate the ability of OL to activate TLR4 and NLRP3 independently of one another. Yet, OL-induced IL-1β release was significantly higher in BMDM from WT than from *Tlr4*^-/-^ mice (Fig. 4 B), indicating that OL signalling through TLR4 contributes to NLRP3 signalling outputs, probably by enhancing the inflammasome priming step.

**Figure 4:**
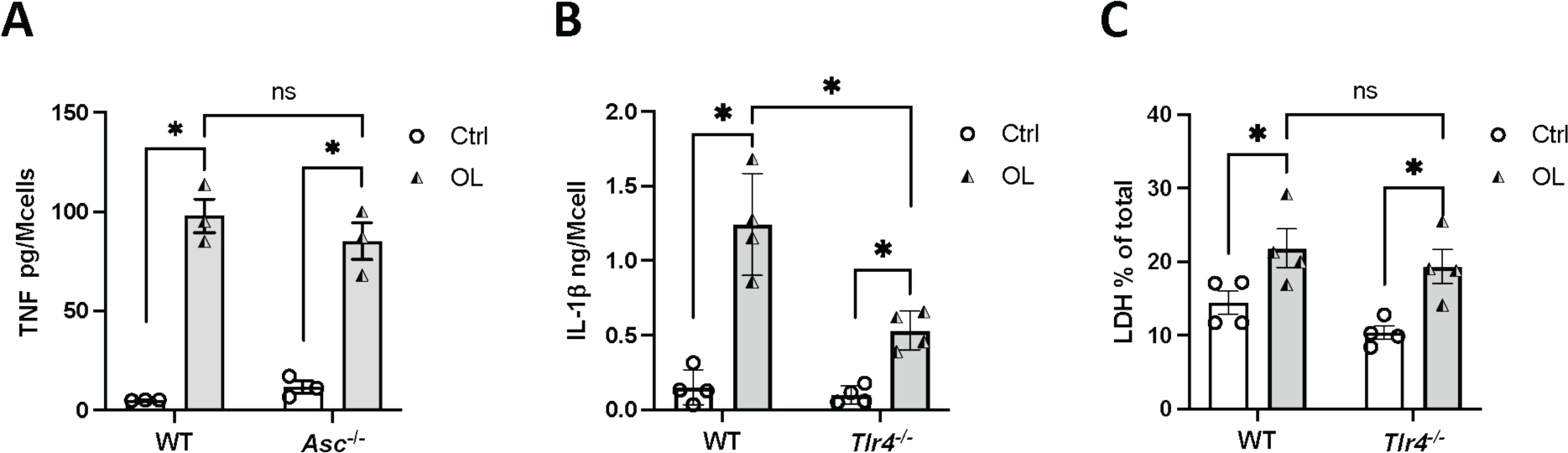
OL ac/vates TLR4 and the inflammasome pathway independently of each other. **A** BMDM from wild type and *Asc*^-/-^ mice were incubated for 4 hours without (Ctrl, white bars) or with OL 100 μM (OL, grey bars) and TNF was quantified in cell supernatants by ELISA. **B and C** BMDM from wild type and *Tlr4*^-/-^ mice were incubated for 4 hours with Pam_3_CSK_4_ (1 μg/mL), then washed and incubated for 18 hours without (Ctrl, white bars) or with OL 100 μM (OL, grey bars). IL-1β was quantified in cell supernatants by ELISA. LDH activity was measured in fresh supernatant by cytotoxicity LDH assay and reported here as the percentage of LDH activity measured in triton-lysed cells. Data information: Bars are the mean of three or four independent experiments represented by symbols (nti 3 to 4) ± SEM. Statistical analysis: Statistical analysis: Unpaired t-test, significant difference for p<0.05 (*). Only comparisons of interest are shown.

### OL primes and activates the NLRP3 inflammasome in murine macrophages and human PBMC

Since OL-induced TLR4 activation enhances OL-induced NLRP3 signalling, we tested whether OL may be capable of both priming and activating NLRP3 in murine and human cells. First, we incubated BMDMs with OL for 4 hours and measured NLRP3 and pro-IL-1β mRNA expression. In line with OL activating TLR4, we found that OL induced NLRP3 and IL-1β mRNA transcription (Fig. 5 A and B). To further confirm this finding, we monitored NLRP3 protein expression in wild-type BMDM and pro-IL-1β protein in *Nlrp3*^-/-^ BMDM, upon cell treatment with OL for 4 hours, using LPS as positive control to upregulate expression of these proteins. As shown in Figures 5 C and D, OL induced NLRP3 and pro-IL-1 β expression, demonstrating the capacity of OL to prime the cells for NLRP3 activation. Together, these data indicate that OL can prime murine macrophages for inflammasome signalling.

**Figure 5:**
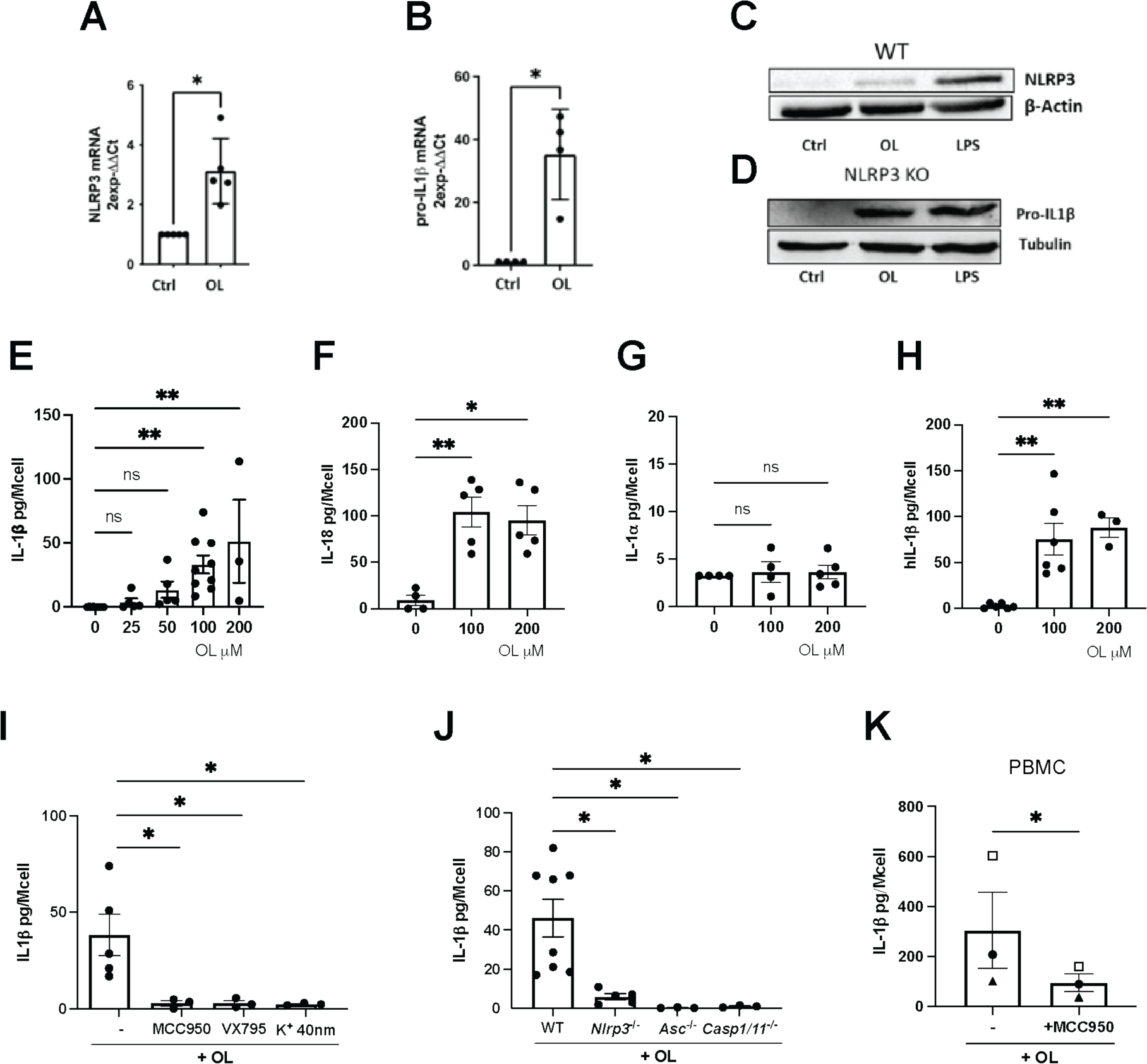
OL primes and ac/vates the canonical NLRP3 inflammasome in both human and murine unprimed cells. **A to B** BMDM from wild-type mice were incubated for 4 hours with medium (Ctrl) or OL 100 μM. NLRP3 (A) or IL-1β (B) mRNA expression was quantified by qPCR and reported here as normalized to untreated cells (Ctrl). **C** BMDM from wild-type mice were incubated for 4 hours with medium (Ctrl), OL 100 μM, or LPS 100 ng/mL, and NLRP3 protein expression was quantified by western blot. **D** BMDM from *Nlrp3*^-/-^mice were incubated for 4 hours with medium (Ctrl), OL 100 μM, or LPS 100 ng/mL, and IL-1β protein expression was quantified by western blot. **E** BMDM from wild-type mice were incubated for 18 hours with medium (0) or with OL 25, 50, 100, or 200 μM, and IL-1β was quantified in cell supernatants by ELISA. **F to G** BMDM from wild-type mice were incubated for 18 hours with medium (0) or with OL 100 or 200 μM, and IL-18 and IL-1⍺ were quantified in cell supernatants by ELISA. **H** Human PBMC from healthy donors were incubated for 18 hours without (0) or with OL 100 or 200 μM, and IL-1β was quantified in cell supernatants by ELISA. **I** BMDM from wild-type mice were incubated for 18 hours with OL 100 μM in the absence (-) or presence of 10 μM NLRP3 inhibitor MCC950, 10 μM of caspase-1 inhibitor VX-765, or in a high K^+^ buffer (KCl 40 mM), and IL-1β was quantified in cell supernatants by ELISA. **J** BMDM from wild-type (WT), *Nlrp3*^-/-^, *Asc*^-/-^, or *Casp1/11*^-/-^ mice were incubated for 18 h with OL 100 μM, and IL-1β was quantified in cell supernatants by ELISA. **K** Human PBMC from healthy donors were incubated for 18 hours with OL 100 μM in the absence or presence of 10 μM MCC950, and IL-1β was quantified in cell supernatants by ELISA. **Data informa/on:** Bars are the mean of at least three independent experiments represented by symbols (n = 3 to 8) ± SEM. **Sta/s/cal analysis: A to B** One-sample t-test. **E to J** Ordinary one-way ANOVA Dunnej’s multiple comparisons test (unpaired). **K** ratio t-test. Significant difference for p<0.05 (*). Only comparisons of interest are shown.

To investigate the ability of OL not only to prime but also activate the inflammasome in unprimed cells, we measured inflammasome-dependent cytokines in supernatants of unprimed BMDMs and PBMC treated with increasing doses of OL for 18 hours. As shown in Figure 5 E, OL induced a weak but dose-dependent IL-1β secretion in BMDMs, significantly higher than untreated conditions when OL was used at high concentrations (100 and 200 µM). Supernatants from BMDM treated with high dose OL also contained significant amounts of IL-18 but not IL-1α (Fig. 5 F and G). OL also induced IL-1β secretion in unprimed human PBMC (Fig. 5 H), indicating that OL activates the inflammasome in unprimed human and murine immune cells.

To further investigate whether OL activates the NLRP3 inflammasome in unprimed cells and whether this was dependent on potassium (K^+^) efflux, we incubated BMDMs with OL in the absence or presence of the NLRP3 inhibitor MCC950, the caspase-1/4 inhibitor VX-765, or a high K^+^ buffer. We found that OL-induced IL-1β secretion was abrogated by NLRP3 and caspase-1/4 inhibitors, and by the high K^+^ buffer (Fig. 5 I). These data thus suggest that OL activates the NLRP3-ASC-casp1 inflammasome in a K^+^ efflux-dependent manner in unprimed cells. To further confirm this finding, we also compared IL-1β secretion induced by OL in BMDM derived from wild-type, NLRP3 (*Nlrp3*^-/-^), ASC (*Asc*^-/-^), or caspase-1/11 (*Casp1/11*^-/-^) mice. As with our findings in primed cells, OL-induced IL-1β required NLRP3, ASC, and casp-1 (Fig. 5 J), confirming that OL activates the NLRP3 inflammasome in unprimed murine macrophages. In further experiments investigating whether OL also activates NLRP3 in human primary cells, we incubated PBMC for 18 hours with OL in the absence or presence of the NLRP3 inhibitor MCC950, and found that MCC950 abrogated OL-induced IL-1β secretion (Fig. 5 K). These data collectively indicate that OL activates the NLRP3 inflammasomes in primary human and murine cells, where OL can provide both the priming and activation signal for the NLRP3 inflammasome.

## Discussion

Previous studies reported on the ability of OL to induce IL-6, TNF, and IL-1β secretion (Kawai and Akagawa, 1989; Okemoto *et al*., 2008), while other studies claimed that OL is either inert or an inhibitor of LPS-induced inflammatory reactions (Kawai *et al*., 1991; Palacios-Chaves *et al*., 2011). Such studies used OL purified from different bacterial strains but also employed different experimental conditions such as cell type, *in vivo* model, incubation time and the presence or absence of other bacterial components such as LPS. Thus, it remained unclear whether: (i) OL is a pro-or anti-inflammatory lipid, (ii) OLs from different bacterial origins exert distinct effects, or (iii) some of the observed pro-inflammatory effects of OL reflected contamination with bacterial LPS. Using synthetic OL free of LPS contamination, we demonstrated that OL is a partial TLR4 agonist that drives context-specific immune reactions, as well as a novel activator of the NLRP3 inflammasome.

OL induces macrophages to secrete TNF. Here, we elucidated the function of TLR4 activation in OL-induced TNF secretion. We discovered that TLR4-blocking antibodies, as well as macrophage TLR4 deficiency, suppressed OL-induced TNF. Thus OL triggers TLR4 signalling to induce the secretion of TNF from macrophages.

TLR4 activation is known to prime inflammasome responses(Schroder and Tschopp, 2010). In the current study, we show that like the canonical TLR4 agonist LPS, OL induced the expression of NLRP3 and pro-IL-1β proteins, priming the macrophages for inflammasome activation. Some NLRP3 priming agents can also provide the activation signal for NLRP3 (e.g., hyaluronan, biglycan, and heme) (Babelova *et al*., 2009; Yamasaki *et al*., 2009; Erdei *et al*., 2018). We discovered that similarly,OL both primes and activates the canonical NLRP3 inflammasome. We showed that OL induced IL-1β secretion in both primed and unprimed human and murine cells, and OL-induced IL-1β secretion was blocked by NLRP3 or casp-1 inhibitors, or cell deficiency in these proteins. These findings explain the previously observed ability of OL to induce both TNF and IL-1β (Kawai and Akagawa, 1989; Okemoto *et al*., 2008). Our data collectively identify OL as both a priming agent and a novel agonist of the NLRP3 inflammasome.

The ability to activate both TLR4 and the NLRP3 inflammasome was also observed for synthetic ionisable lipids used as transfection agents (Lonez *et al*., 2014; Pizzuto *et al*., 2018), but the physiological relevance of such activities was unclear as the capacity of analogous lipids found in nature was untested. Our study demonstrates the capacity of a bacterial ionisable cationic lipid to modulate the immune system through TLR4 and NLRP3. It is likely that OL shares with cationic lipids the capacity to activate TLR4 TRIF and MyD88 signalling pathways, as well as inducing lysosomal destabilisation and cathepsin release for NLRP3 activation (Li *et al*., 2018; Alarcón-Vila, Pizzuto and Pelegrín, 2019; He *et al*., 2019; Pizzuto, Pelegrin and Ruysschaert, 2022). Thanks to the ability of ionisable cationic lipids to deliver antigens while activating TLRs and NLRP3 and stimulating the adaptative immune response required for successful vaccine formulation (Demento *et al*., 2009; Awate, Babiuk and Mutwiri, 2013; Pulendran, S. Arunachalam and O’Hagan, 2021; Tahtinen *et al*., 2022), treatment of mice with an antigen simply mixed with the lipids resulted in effective, adjuvant-free vaccination (Pizzuto *et al*., 2018). OL’s ability to transport antigens or deliver nucleic acids while activating TLR4 and NLRP3 is worth exploring to support its use as an adjuvanted carrier in new vaccine technologies.

Previous studies of synthetic ionisable cationic lipids did not determine whether these lipids could stimulate NLRP3 versus TLR4 independently of one another. Here, we show that OL activation of TLR4 and NLRP3 can be uncoupled. We show that OL-induced NLRP3 activation and resultant IL-1β and LDH release can occur independently of TLR4 activation (*Tlr4*-/- BMDM) if the priming of NLRP3 was achieved using the TLR2 activator, Pam_3_CSK_4_. A similar uncoupling of NLRP3 from TLR4 signaling was previously observed for LPS *in vitro* and *in vivo* (Kayagaki *et al*., 2013), with LPS activating TLR4-independent NLRP3 downstream of the non-canonical inflammasome when Pam_3_CSK_4_ or poly(I:C) are used as priming agents (Kayagaki *et al*., 2013). Likewise, OL-induced TLR4 signaling may be uncoupled from NLRP3 signaling, with TLR4 activation by OL observed here in inflammasome-deficient murine macrophages, as expected for a TLR4 agonist (Gay *et al*., 2014). It bears noting that in human monocytes, LPS-induced TLR4 activation is sufficient to activate NLRP3 through its alternative pathway(Gaidt *et al*., 2016). Therefore, further investigation is required to elucidate the TLR4 dependency of OL-induced NLRP3 signalling in human monocytes.

While OL, like LPS, can activate TLR4 and NLRP3 independently in macrophages, it is a less potent inducer of cytokine production compared to LPS. Some weak receptor agonists are defined as partial agonists if they can block the response of stronger receptor ligands(Kenakin, 2017). Here, we discovered that OL suppresses LPS-induced TLR4 signalling, and is thus a partial agonist of TLR4. This suppressive effect of OL was specific to TLR4, as OL did not suppress cell responses to TLR2 and NLRP3 activators. By performing dose titrations of LPS in TLR4-expressing HEK293T cells in the absence or presence of OL, we demonstrated that OL is a partial agonist of TLR4, which might explain the apparently contradictory activities of OL reported in the literature (Kawai and Akagawa, 1989; Kawai *et al*., 1991; Okemoto *et al*., 2008; Palacios-Chaves *et al*., 2011). The dose titrations of LPS presented here show that in the presence of OL, LPS reached the same maximal response of NF-κB activation as in the absence of OL, although higher doses of LPS were needed. This may indicate that OL inhibits TLR4 by binding to the same binding site as LPS. Of note, this highlights a different mechanism of TLR4 interaction with ionisable lipids compared to permanently charged cationic lipids that bind TLR4 at a distinct location distal to the LPS-binding region of TLR4 (Lonez *et al*., 2015).

We also discovered that OL suppressed IL-1β secretion induced downstream of the non-canonical inflammasome in murine macrophages transfected with LPS. This was due to OL inhibiting TLR4-dependent LPS-induced inflammasome priming, rather than inflammasome activation, because this suppressive effect of OL was absent in *Tlr4-*deficient macrophages.

Our identification of OL as an inhibitor of LPS-TLR4 signalling provides a molecular mechanism for a previous phenomenological report that OL inhibits *in vivo* LPS responses such as TNF secretion and LPS-induced lethality (Kawai *et al*., 1991). It also highlights that OL, as an inhibitor of inflammasome priming but not activation, may not be sufficient to protect from LPS-induced lethality in the context of a bacterial infection or sepsis where other priming agents are present.

The capacity of OL to suppress LPS-TLR4 signalling has some similarities to anti-inflammatory LPS homologs produced by Gram-negative bacteria that suppress LPS-TLR4 signalling to avoid immune system detection, and thereby promote pathogenicity (Matsuura, 2013). It is thus tempting to speculate that for bacteria that constitutively produce both OL and LPS, OL may prevent LPS recognition by TLR4 as a microbial evasion strategy to escape host immune surveillance.

On the other hand, the ability of OL to induce TLR4 and NLRP3 signaling has similarities to pro-inflammatory LPS produced by Gram-negative bacteria and detected by our immune system as a signal of a bacterial threat (Aachoui *et al*., 2013; Zanoni and Granucci, 2013; Shi *et al*., 2014; Baker *et al*., 2015). Therefore, it is worth considering that in the context of LPS-free bacteria (e.g., Gram-positive bacteria; Gram-negative bacteria grown in phosphate-depleted environments), the host responds by detecting OL through the TLR4 and the NLRP3 inflammasome pathways, prompting the initiation of an antibacterial immune response.

Although we showed that, in the absence of other stimuli, OL is a weak inducer of cytokine secretion, the activation of the TLR4 and the NLRP3 inflammasome pathways might be more sustained in the context of a bacterial infection where other TLR activators are present. Indeed, we observed that the IL-1β secretion induced by OL in macrophages primed with the TLR2 activator Pam_3_CSK_4_ was 10-30 times higher than that induced in unprimed cells, in line with previous reports showing that the combination of OL with other signals enhances IL-1β secretion (Kawai and Akagawa, 1989). Moreover, bacteria may produce OL species with different chain lengths and saturations from the one tested in this study; those bacterial species might present a stronger immunogenicity, as chain lengths and saturations of lipids have been shown to strongly affect their potency as NLRP3 and TLR4 activators (Li *et al*., 2018; Pizzuto *et al*., 2018, 2019; Alarcón-Vila, Pizzuto and Pelegrín, 2019; He *et al*., 2019; Pizzuto, Pelegrin and Ruysschaert, 2022).

In conclusion, our findings identify OL as a bacterial ionisable lipid able to modulate both TLR4 and NLRP3. Synthetic ionisable lipids have been used as transfection agents for decades. Thus, OL’s potential for antigen transportation or nucleic acid delivery merits investigation to strengthen its application as an adjuvanted carrier in novel vaccine technologies. Moreover, the activation of TLR4 and NLRP3 pathways by bacterial OL might be responsible for pro-inflammatory reactions in the host and be used by our immune system to detect Gram-positive bacteria, or Gram-negative bacteria growing in low-phosphate environments where LPS is not produced. Conversely, when OL is constitutively produced together with LPS, OL may inhibit LPS-TLR4 signalling to allow bacteria to escape host immune surveillance. The role of OL in bacterial immunogenicity or escape strategies may depend on the bacteria species and should be addressed in future investigations.

## Authors’ Contributions

Conceptualisation MP and J-MR. Methodology Investigation MP, LH-N, CM-L JS, MG, and SR-L. Validation MP. Formal analysis MP. Resources KS and PP. Writing-original draft MP. Writing-Review and Editing MP, J-MR, PP and KS. Visualisation MP. Supervision MP, J-MR, KS and PP.

## Acknowledgments

We gratefully acknowledge Dr. Fiona Wylie and Dr. Madhavi Maddugoda for editing this manuscript. We thank Maria de Carmen Baños and Ana Isabel Gomez (IMIB, Murcia, Spain) for technical assistance with cell differentiation and culture. We acknowledge the use of Grammarly and the language model provided by OpenAI for assistance in proofreading, grammar checking, and identifying synonyms during the preparation of this manuscript.

