## Supplementary data for "Ornithine Lipid is a Partial TLR4 Agonist and NLRP3 Activator"

**Expanded View File for:**

Pizzuto et al., 2024

**Content:**

- Expanded data
- Expanded abbreviations
- Expanded references

**Expanded data**

**Synthesis of Ornithine Lipid**

**Chemistry**

To obtain the ornithine lipid compound, an amide bond coupling ornithine residue and fatty acyl group was performed via several steps. Briefly, the commercially available α-Fmoc-δ-Boc-L-Orn-OH (compound **1**) was benzylated by Benzyl alcohol after the activation of carboxyl group of ornithine derivative by dicyclohexylcarbodiimide (DCC) and 4-dimethylaminopyridine (DMAP) to obtain compound **2** in good yield (70 %) (scheme 1) (Loke *et al.*, 2012). The N α -Fmoc group was then cleaved to give compound **3** in 84% yield (scheme 1) (Larionov OV, 2004).

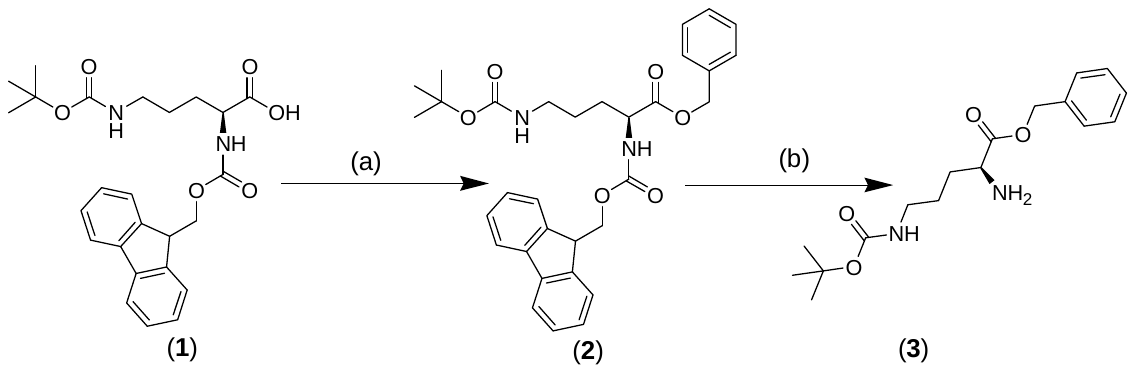

**Scheme 1:** ***Reagents and conditions:*** (a) DCC, DMAP, CH_2_Cl_2_, 0 °C, 24h; (b) pyrrolidine, CH_2_Cl_2_, RT, 1h.

(R,S) 3-hydroxytetradecanoic acid (compound **4**) was converted into its benzyl ester (compound **5**) in quantitative yield by reaction with benzyl bromide, triethylamine (TEA), and tetrabutylammonium iodide (Bu_4_NI). Then the obtained benzylated compound was esterified with myristoyl chloride to give compound **6**, which was purified and submitted to hydrogenation by Pd on carbon to give the fatty acyl moiety (compound **7**) in 99% yield (scheme 2)[3].

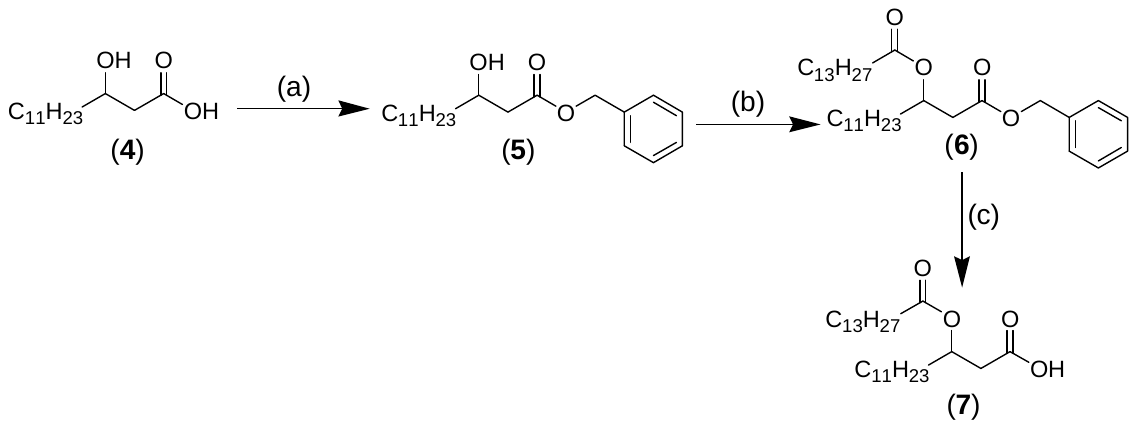

**Scheme 2:** ***Reagents and conditions:*** (a) benzyl bromide, Bu_4_NI, TEA, EtOAc, RT, 24h; (b) myristoyl chloride, pyridine, CH_2_Cl_2_, 0 °C then RT; (c) Pd/C, H_2_, RT, 3h.

The amide bond between ornithine and fatty acyl group was achieved by the acylation of the free amino group of compound **3** with compound **7**, in the presence of isobutyl chloroformate (i-BuOCOCl) and N-methylmorpholine, to give the lipid ornithine compound (**8**) in 67% yield. The benzyl and the δ-Boc groups were cleaved respectively to give compound **10** in excellent yield (scheme 3). Because the fatty acyl residue was obtained starting from R and S 3-hydroxytetradecanoic acid, two isomers of compound **10** were obtained RS and SS [3].

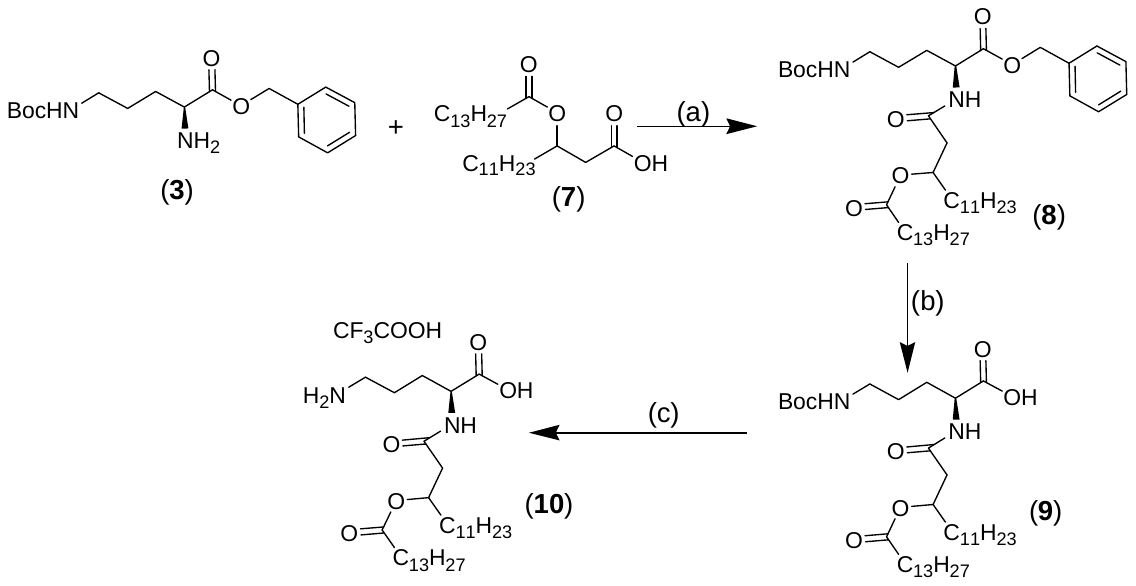

**Scheme 3:** ***Reagents and conditions:*** (a) i-BuOCOCl, N-methylmorpholine, THF, 30 min at -15 °C then RT 24h; (b) Pd/C, H_2_, RT, 2h; (c) CF_3_COOH.

**Materials**

^1^H NMR spectra were taken on a Bruker Avance 400 M Hz spectrometer (Wissemburg, France) at 293 K. Chemical shifts (δ) are given in parts per million (ppm), and the coupling constants are expressed in hertz. Mass spectrometric data were obtained on a QTOF 6520 (Agilent, Palo Alto, CA), positive mode, electrospray ionization (ESI), mode time of flight (TOF), by diffusion of 0.5 mL/min, by mobile phase CH_3_OH (VCAP 3500 V, source *T*, 350 °C; fragmentation, 110 V; skimmer, 65 V). All reactions were followed by thin-layer chromatography (TLC) carried out on Fluka (Bornem, Belgium) PET foils silica gel 60, and compounds were visualized by UV and by spraying with 5% p-anisaldehyde in methanol. Column chromatographies were performed with EchoChrom MP silica 63–200 from MP Biomedicals (Santa Ana, CA). Organic solutions were dried over Na_2_SO_4_ and concentrated with a Buchi rotatory evaporator (Flawil, Switzerland). Starting materials 3-hydroxymyristic acid (racemic), myristoyl chloride , N-alpha-(9-fluorenylmethyloxycarbonyl)-N-delta-t-butyl-oxycarbonyl-L-ornithine , benzyl alcohol , benzyl bromide , tetrabutylammonium Iodide , and N,N'-dicyclohexylcarbodiimide (DCC) were available from TCI (Japan). Isobutyl chloroformate and 4-methylmorpholine were purchased from Sigma-Aldrich (Bornem, Belgium).

**Synthesis**

**Nδ -tert-Butoxycarbonyl-Nα -fluoren-9-ylmethoxycarbonyl-L-ornithine benzyl ester (2)** (Loke *et al.*, 2012)

Benzyl alcohol (1 g, 9.2 mmol) and 4-dimethylaminopyridine (0.1 g, 0.8 mmol) were added to a solution of Fmoc-Orn(Boc)-OH (compound **1**) (3 g, 6.6 mmol) in CH_2_Cl_2_ (100 mL) and the mixture was cooled in an ice bath. Dicyclohexylcarbodiimide (1.5 g, 7.2 mmol) was added and stirring was continued overnight, allowing the mixture to warm up to RT. The white solid was removed by vacuum filtration. The filtrate was washed with 2% KHSO_4_ in water (50 mL), 5% KHCO_3_ in water (50 mL), water (30 mL). After drying over Na_2_SO_4_, the solvent was removed and the residue subjected to column chromatography (eluent: CH_2_Cl_2_ then CH_2_Cl_2_/EtOAc 8:1) (R_f_ 0.5). Removal of the solvent from the eluate afforded the product as a white solid (2,5 g, 70%) mp 101−104 °C. ^1^H-NMR (400 MHz, CDCl_3_): δ (ppm) 7.69 (d, *J=* 8 Hz, 2H) 7.52 (d, *J=* 8 Hz, 2H), 7.28−7.41 (m, 9H), 5.40 (d, *J=* 8 Hz, 1H), 5.09 (dd, *J*= 12, 8 Hz, 2H), 4.63 (d, *J=* 4 Hz, 1H), 4.43 (m 1H), 4.33 (d, *J*= 4 Hz, 2H), 4.13 (m, 1H), 3.03 (br s, 2H), 1.80 (m, 1H), 1.58 (m, 1H), 1.36−1.50 (m, 11H).

**(S)-Benzyl 2-amino-5-((tert-butoxycarbonyl)amino)pentanoate (3)** (Larionov OV, 2004)

To a solution of compound **2** (2 g, 3,5 mmol) in 10 mL CH_2_Cl_2_ was added 10% v/v pyrrolidine in CH_2_Cl_2_ (10 mL) and stirred for 1h. Then the solvent was evaporated and the crude product purified with column chromatography (silica gel, hexane/EtOAc 1 : 5 then CH_2_Cl_2_/CH_3_OH 9:1) to give compound **3** (1g, 85%) as an yellow oil. R_f_ = 0.1 (CH_2_Cl_2_/CH_3_OH 9:1, the primary amine of compound **3** was detected by a solution of 5% *p*-anisaldehyde in methanol). ^1^H-NMR (400 MHz, CDCl_3_): δ (ppm ): 7.35 (m, 5H, Ar), 5.15 (s, 2H, CH_2_Ph), 4.63 (m, 1H, NH), 3.48 (m, 1H, 2-H), 3.11 (m, 2H, 5-H), 1.75 (m, 1H, H-3), 1.56 - 1.6 (m, 5H, 4-H, 3-H, NH_2_), 1.40 (s, 9H, C(CH_3_)_3_).

**Benzyl 3-hydroxytetradecanoate (5)** (Martin *et al.*, 2006)

To a suspension of 3-hydroxytetradecanoic acid (compound **4**) (2g, 8.2 mmol) in ethyl acetate (40 mL) were added benzyl bromide (3 ml, 25 mmol), triethylamine (3.4 ml, 24 mmol), and tetrabutylammonium iodide (1.6 g, 4.3 mmol). The mixture was stirred for 18 h at room temperature. The solvent was then evaporated, and the residue was taken up in ether (40 mL) and washed with saturated aqueous NaHCO_3_ (50 mL) and then with water (50 mL). The organic phase was separated and dried over Na_2_SO_4_. The solvent was evaporated, and the compound was purified by column chromatography (silica gel, CH_2_Cl_2_/EtOAc 9:1, *R_f_*= 0.2) thus affording 1.9 g (71% yield) benzyl 3-hydroxytetradecanoate **3.** ^1^H-NMR (400 MHz, CDCl_3_): δ (ppm ): 7.36 (m, 5H, Ar), 5.16 (s, 2H, *CH_2_*Ph), 4.02 (m, 1H, H-3), 2.85 (d, *J*= 4 Hz, 1H, -OH), 2.56 (dd, *J*= 4, 16 Hz, 1H, H-2A), 2.46 (dd, *J*= 8, 16 Hz, 1H, H-2B), 1.56-1.26 (m, 20H, CH_2_), 0.88 (t, *J*= 8 Hz, CH_3_).

**Benzyl 3-(tetradecanoyloxy)tetradecanoate (6)** (Martin *et al.*, 2006)

Compound **5** (1 g, 2.99 mmol) was acylated with myristoyl chloride (0.9 mL, 3.32 mmol) in a mixture of pyridine (1 mL) and CH_2_Cl_2_ (50 mL) at 0 °C and then at room temperature for 24 h. The reaction mixture was poured into ice−water containing 5% aqueous NaHCO_3_ and the organic phase was separated. The organic phase was washed with 1 N aqueous HCl and then with brine (50 mL) and water (50 mL). The organic phase was separated, dried over Na_2_SO_4_, and filtered. The solvent was evaporated and the residue was purified by column chromatography on silica gel (CH_2_Cl_2_/hexan, 1:1) to give **6** (1.1 g, 67% yield)  as a colorless syrup then white wax:  ^1^H NMR (400 MHz, CDCl_3_) δ 7.35-7.25 (m, 5H, Ph), 5.22 (m, 1H, H-3), 5.10 (s, 2H, C*H_2_*Ph), 2.59 (m, 2H, H-2B and H-2A), 2.18 (m, 2H,  H-2’), 1.54 (m, 4H, H-4, H3’), 1.29−1.24 (br s, 38H, 19 CH_2_), 0.87 (t, 6H, *J*= 8 Hz, 2 CH_3_).

**3-(tetradecanoyloxy)tetradecanoic acid (7)**

To a solution of benzyl ester **6** (1.1 g, 2 mmol) in a mixture of ethyl acetate (30 mL) and methanol (30 mL) was added 10% Pd on carbon (300 mg). The mixture was hydrogenated at room temperature under atmospheric pressure of hydrogen for 1 h. The catalyst was then removed by filtration on celite and the solvent was evaporated to give pure **7**  (0.9 g, 99%) as a colorless syrup then white wax:  ^1^H NMR (400 MHz, CDCl_3_) δ 5.21 (m, 1H, H-3), 3.54 (br s, 1H, -OH), 2.63-2.52 (m, 2H, H-2B and H-2A), 2.27 (t, *J=* 8 Hz, 2H,  H-2’), 1.59 (m, 4H, H-3’ and H-4), 1.29−1.24 (br s, 38H, 19 CH_2_), 0.88 (t, 6H, J= 8 Hz, 2 CH_3_).

**Benzyl (2S)-5-((tert-butoxycarbonyl)amino)-2-(3 (tetradecanoyloxy) tetradecanamido) pentanoate (8)** (Martin *et al.*, 2006)

 To a stirred solution of 3-(tetradecanoyloxy)tetradecanoic acid **7**(0.56 g, 1.2 mmol) in THF (15 mL) at −15 °C were added successively *N*-methylmorpholine (0.14 mL, 1.2 mmol) and isobutyl chloroformate (0.16 mL, 1.2 mmol). Stirring was continued for 30 min at −15 °C. A solution of **3** (0.4 g, 1.2 mmol) in THF (15 mL) was then added to the reaction mixture. After stirring overnight at room temperature, the solvent was removed in vacuo and H_2_O (20 mL) was added to the residue. The mixture was then extracted with EtOAc (2 × 30 mL), and the organic phases were combined, washed successively with H_2_O (20 mL) and brine (20 mL), and dried over Na_2_SO_4_. The solvent was evaporated and the residue was purified by column chromatography on silica gel (CH_2_Cl_2_/EtOAc, 8:2) to give **8** (0.6 g, 67%) as a white solid:  *R_f_* = 0.29 (hexane/EtOAc, 1:1); ^1^H NMR (400 MHz, CDCl_3_) δ 7.34−7.36 (m, 5H, Ph), 6.41 (m, 1H, NH), 5.17−5.12 (m, 3H, H-3‘, C*H_2_*Ph), 4.65-4.56 (m, 2H, H-2 and NH), 3.08 (m, 2H, H-5), 2.48 (m, 2H, H-2’), 2.28 (m, 2H, H-2”), 1.84-1.60 (m, 8H, H-3, 4, 3”, 4’), 1.42 (s, 9H, tBu) 1.25 (br s, 38H, 19 CH_2_), 0.88 (t, 6H, *J*= 8 Hz, 2 CH_3_).

**(2S)-5-((tert-butoxycarbonyl)amino)-2-(3-(tetradecanoyloxy)tetradecanamido)pentanoic acid (9)** (Martin *et al.*, 2006)

To a solution of benzyl ester (**8**) (0.6 g, 0.8 mmol) in ethanol (30 mL) was added 10% Pd on carbon (90 mg). The mixture was hydrogenated at room temperature under atmospheric pressure of hydrogen for 2 h. The catalyst was then removed by filtration and the solvent was evaporated to give pure **9** (0.45 g, 88% yield). ^1^H NMR (400 MHz, CDCl_3_) δ 6.87 (m, 1H, NH), 5.18 (m, 1H, H-3’), 4.88 (m, 1H, NH), 4.56 (m, 1H, H-2), 4.19 (br s, 1H, OH), 3.14 (m, 2H, H-5), 2.51 (m, 2H, H-2’), 2.29 (m, 2H, H-2”), 1.91-1.52 (m, 7H, H-3, 4, 3”, 4’), 1.44 (s, 9H, tBu) 1.25 (br s, 38H, 19 CH_2_), 0.88 (t, 6H, *J*= 8 Hz, 2 CH_3_).

**(2S)-5-amino-2-(3-(tetradecanoyloxy)tetradecanamido)pentanoic acid trifluroacetic salt (10)** (Martin *et al.*, 2006)

Compound **9** (0.40 g, 0.9 mmol) was dissolved in trifluoroacetic acid TFA (2 mL) and the solution was stirred at room temperature for 2.5 h. The solvent was then removed under high vacuum to give the desired compound **10** as yellow semisolid (0.4 g, 98% yield). ^1^H NMR (400 MHz, CDCl_3_) δ 6.22 (br s, 2H, COOH, NH), 7.58 (br s, 3H, NH_3_^+^), 5.19 (m, 1H, H-3’), 4.36 (m, 1H, H-2), 3.02 (m, 2H, H-5), 2.52 (m, 2H, H-2’), 2.25 (m, 2H, H-2”), 1.90-1.56 (m, 8H, H-3, 4, 3”, 4’), 1.25 (br s, 38H, 19 CH_2_), 0.88 (t, 6H, *J*= 8 Hz, 2 CH_3_). FTIR: 3334, 3071, 2916, 2850, 1704, 1643, 1537, 1377, 1251, 1197, 1116, 1111, 828, 719 cm^-1^. HRMS (ESI) calculated for C_33_H_64_N_2_O_5_ (M+): 568.4835, found: 568.4842, error: 1.23 ppm. (Fig. S6)

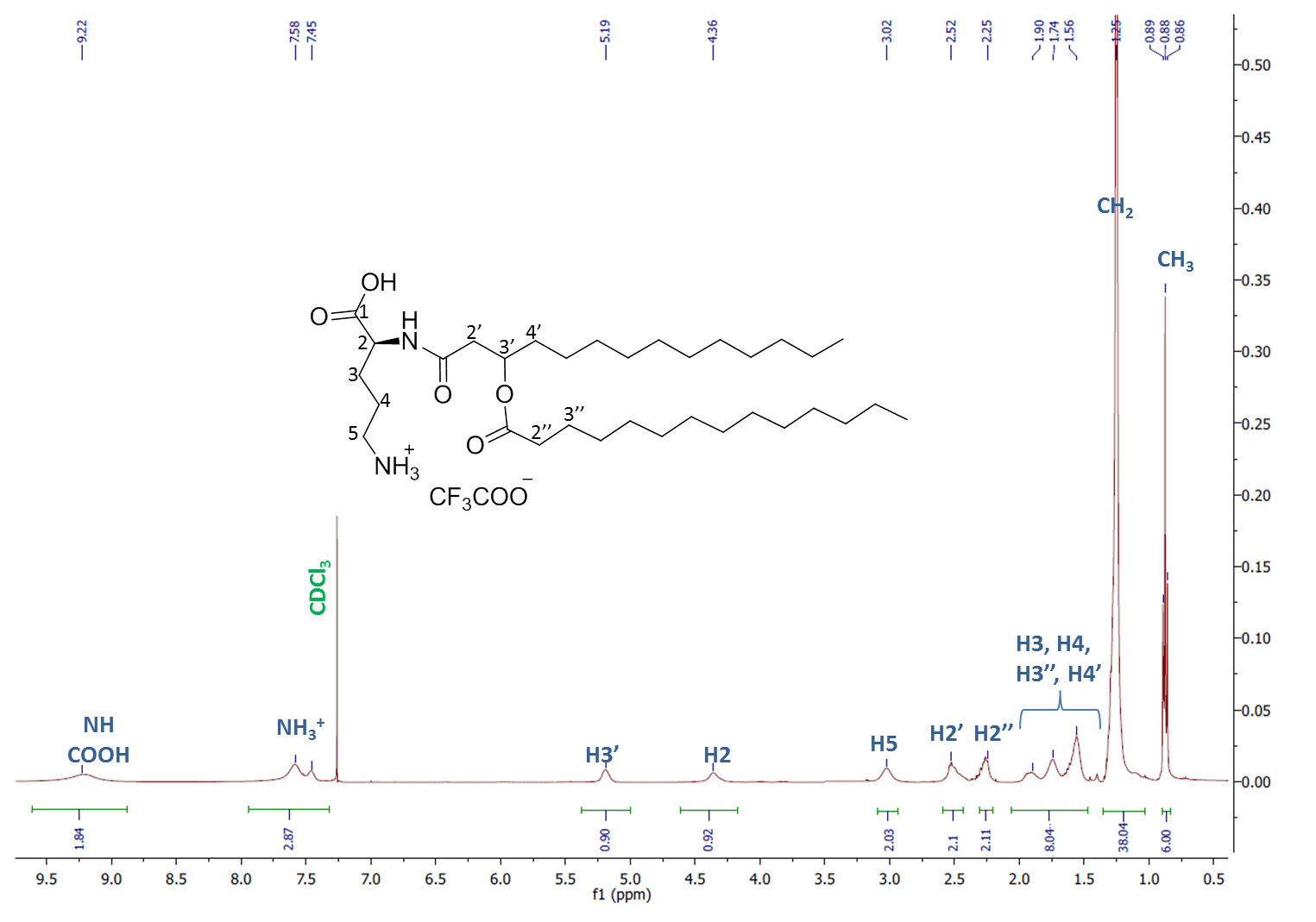

**Expanded figure EV1:** Structure and ^1^H-NMR of compound **10** (Ornithine Lipid)

2. **OL-induced TLR4 activation is not due to LPS contamination.**

To test whether the OL preparation was free of LPS contamination, we used the Chromogenic Endotoxin Quantitation Kit from Thermo Fisher, which allows the quantification of LPS content from 0.01 to 0.1 ng/mL. We quantified LPS content in concentrated OL preparations (334 μM) in the absence or presence of triton 0.05%. Such OL concentrations were more than three times greater than the ones that showed significant TNF secretion in BMDM. The detergent (triton) was added to dissolve OL aggregate and potentially release LPS, if any. The measured LPS content was under the lowest LPS standard and not significantly higher than the blank in both detergent and detergent-free preparations. Therefore, we concluded that LPS content in OL preparation is null. To remove any doubt that minimal LPS contamination under the kit detection limit may account for OL TLR4 activity, we tested whether such LPS content is able to induce TNF secretion in BMDM. We incubated BMDM with increasing LPS concentrations from 0.01 to 200 ng/mL and measured TNF secretion. The lowest LPS standard concentration of 0.01 ng/mL did not induce TNF secretion. Therefore, even if the LPS content in OL preparations was higher than zero, it cannot be responsible for the observed activation of TLR4 by OL.

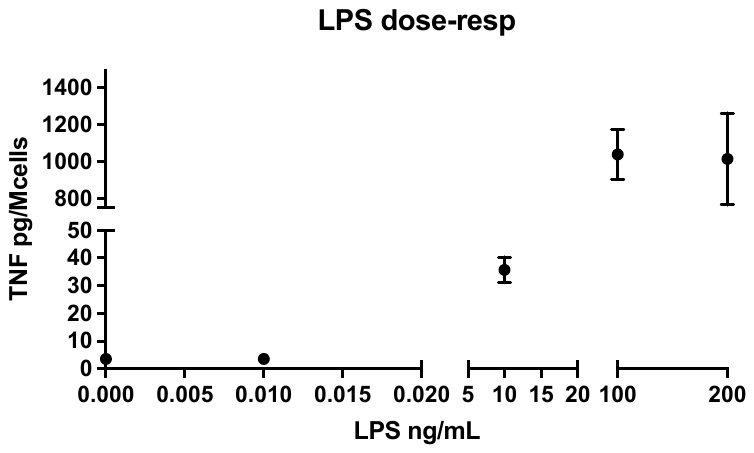

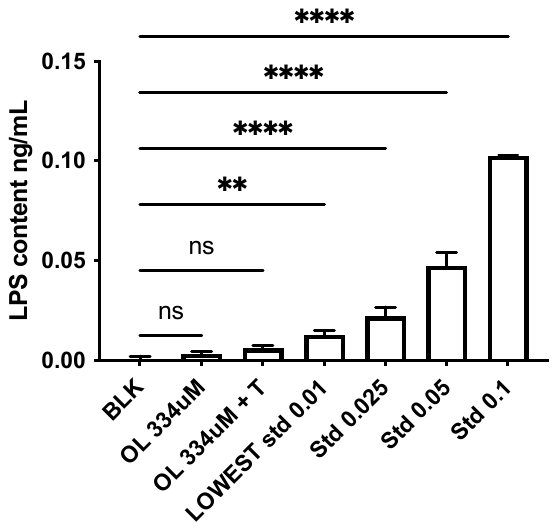

**A**

**B**

**Expanded View Figure EV2:** **OL preparations are free of LPS contamination.**

**A** LPS content was quantified in OL preparations starting stock 334 μM in the absence or presence of 0.05% triton by Chromogenic Endotoxin Quantitation Kit and reported here together with the value measured in LPS preparations used as standard.

**B** Bone marrow-derived macrophages (BMDM) from wild-type mice were incubated for 4 h with increasing LPS concentrations (0.01 to 200 ng/mL). TNF was quantified in collected supernatants by ELISA.

**Data information:** Bars are the mean of three independent experiments represented by symbols (n= 3) ± SEM. In B, symbols represent the mean of four independent experiments (n= 4) ± SEM

**Statistical analysis:** Ordinary ne-way ANOVA Dunnett’s multiple comparisons test (unpaired), significant difference for p<0.05 (*). Only comparison of interest are shown.

**Expanded abbreviations**

dicyclohexylcarbodiimide (DCC); 4-dimethylaminopyridine (DMAP); triethylamine (TEA); tetrabutylammonium iodide (Bu_4_NI); Ethyl acetate (EtOAc); Room temperature (RT); Tetrahydrofuran (THF).

**Expanded References**

Larionov OV, de M.A. (2004) “Enantioselective total syntheses of belactosin A, belactosin C, and its homoanalogue. ,” *Org Lett.*, 6(13), pp. 2153–2156.

Loke, I. *et al.* (2012) “Influence of steric parameters on the synthesis of tetramates from α-amino-β-alkoxy-esters and Ph3PCCO,” *Tetrahedron*, 68(2), pp. 697–704. doi:10.1016/J.TET.2011.10.099.

Martin, O.R. *et al.* (2006) “Synthesis and immunobiological activity of an original series of acyclic lipid a mimics based on a pseudodipeptide backbone,” *Journal of medicinal chemistry*, 49(20), pp. 6000–6014. doi:10.1021/JM060482A.
